# Overloading And unpacKing (OAK) - droplet-based combinatorial indexing for ultra-high throughput single-cell multiomic profiling

**DOI:** 10.1101/2024.01.23.576918

**Authors:** Bing Wu, Hayley M. Bennett, Xin Ye, Akshayalakshmi Sridhar, Celine Eidenschenk, Christine Everett, Evgeniya V. Nazarova, Hsu-Hsin Chen, Ivana K. Kim, Margaret Deangelis, Leah A. Owen, Cynthia Chen, Julia Lau, Minyi Shi, Jessica M. Lund, Ana Xavier-Magalhaes, Neha Patel, Yuxin Liang, Zora Modrusan, Spyros Darmanis

## Abstract

Multiomic profiling of single cells by sequencing is a powerful technique for investigating cellular diversity in complex biological systems. Although the existing droplet-based microfluidic methods have advanced single-cell sequencing, they produce a plethora of cell-free droplets and underutilize barcoding capacities due to their low cell concentration prerequisites. Meanwhile, combinatorial indexing on microplates can index cells in a more effective way; however, it requires time-consuming and laborious protocols involving multiple splitting and pooling steps. Addressing these constraints, we have developed “Overloading And unpacKing” (OAK). With reduced labor intensity, OAK can provide cost-effective multiomic profiling for hundreds of thousands of cells, offering detection sensitivity on par with commercial droplet-based methods. To demonstrate OAK’s versatility, we conducted single-cell RNA sequencing (scRNA-Seq) as well as joint single-nucleus RNA sequencing (snRNA-Seq) and single-nucleus Assay for Transposase Accessible Chromatin with sequencing (snATAC-Seq) using cell lines. We further showcased OAK’s performance on more complex samples, including *in vitro* differentiated bronchial epithelial cells and primary retinal tissues. Finally, we examined transcriptomic responses of 408,000 melanoma cells across around 1,000 starting lineages over a 90-day treatment with a RAF inhibitor, belvarafenib. We discovered a rare cell population (0.12%) that underwent a sequence of transcriptomic changes, resulting in belvarafenib resistance. Ultra-high throughput, broad compatibility with diverse molecular modalities, high detection sensitivity, and simplified experimental procedures distinguish OAK from previous methods, and render OAK a powerful tool for large-scale analysis of molecular signatures, even for rare cells.

## Main

The technological landscape of single-cell sequencing is rapidly evolving, encompassing newly developed methods^1–4^ that offer an unprecedented view of cellular heterogeneity. This technical evolution is fueled by the need to achieve more precise cell type or state identification, capture rare cell states or cellular lineages, and conduct comprehensive perturbation screens for new drug target discovery, all of which have steered technological development toward analyzing a greater number of cells at a reduced cost.

Droplet-based microfluidic approaches co-encapsulate a barcoded bead and a cell within an emulsion to enable parallel analysis of thousands of individual cells^5–7^. These methods have been an important advancement in streamlining high-throughput single cell sequencing.

However, the low cell concentration required to minimize the number of multi-cell droplets leads to a large number of cell-free droplets and underutilized barcoding capacity. Alternatively, combinatorial indexing on microwell plates^8,9^ provides a strategy for barcoding over 100,000 cells^10–12^. However, this ultra-high throughput approach comes with long and laborious protocols, involving multiple rounds of splitting and pooling cells for indexing.

Inspired by the strengths and limitations of these two families of single cell sequencing methods, we have developed OAK, a novel approach that combines droplets with combinatorial indexing to achieve both elevated throughput and experimental simplicity. OAK can be used to measure gene expression, accessible chromatin, and antibody conjugated oligonucleotides, either separately or jointly. With OAK, we performed paired snRNA-Seq and snATAC-Seq on complex retinal tissue. Furthermore, we undertook a lineage tracing experiment capturing RNA and lineage barcodes for 408,000 cells, revealing the longitudinal response of melanoma cells to a RAF inhibitor, belvarafenib. Compatibility with diverse molecular modalities, high detection sensitivity, and easier experimental procedures distinguish OAK from previous methods^13–15^ that conduct combinatorial indexing with the use of microfluidics.

## Results

### Principles and performance of OAK

OAK relies on fixed cells or nuclei to serve as individualized reaction chambers for two rounds of indexing (Fig. 1a). The first round is conducted within droplets, for which we utilized a commercially available system, the Chromium system by 10x Genomics. In this and other droplet-based single-cell sequencing systems^5–7,16^, the cells are loaded at a low concentration to minimize the possibility of encapsulating multiple cells within a single droplet. Based on the Poisson distribution, this is estimated to result in over 80% of droplets devoid of a cell (Fig. 1b), leaving their barcoding potential untapped and reagent wasted. To more efficiently utilize the droplets, we overloaded the microfluidic chip in the Chromium system, resulting in reduced cell-free droplets, concomitant with increased single- and multi-cell droplets (Fig. 1b). To resolve single cells in multi-cell droplets, after the first round of indexing mediated by in-situ reverse transcription of mRNA, we unpacked droplets by breaking emulsions (Fig. 1a). Thus, all encapsulated cells are released, mixed, and randomly distributed into multiple aliquots. The number of aliquots to generate can be tuned based on the scale of cell loading and the number of droplets made by the microfluidic system, in order to achieve a desirable theoretical multiplet rate (Extended Data Fig. 1a). Each aliquot will receive a unique secondary index integrated to molecules that already carry primary indexes (Fig. 1a). From this secondary indexing step, researchers can select any scale of cell subsets to create sub-libraries for sequencing, a feature only shared by select droplet-based methods^5^. This enables an increasing number of cells to be sequenced in a stepwise manner to assess data quality, to receive sufficient sequencing depth, or to avoid sequencing excessive cells, which will be further described in the subsequent sections of this study.

**Figure 1:**
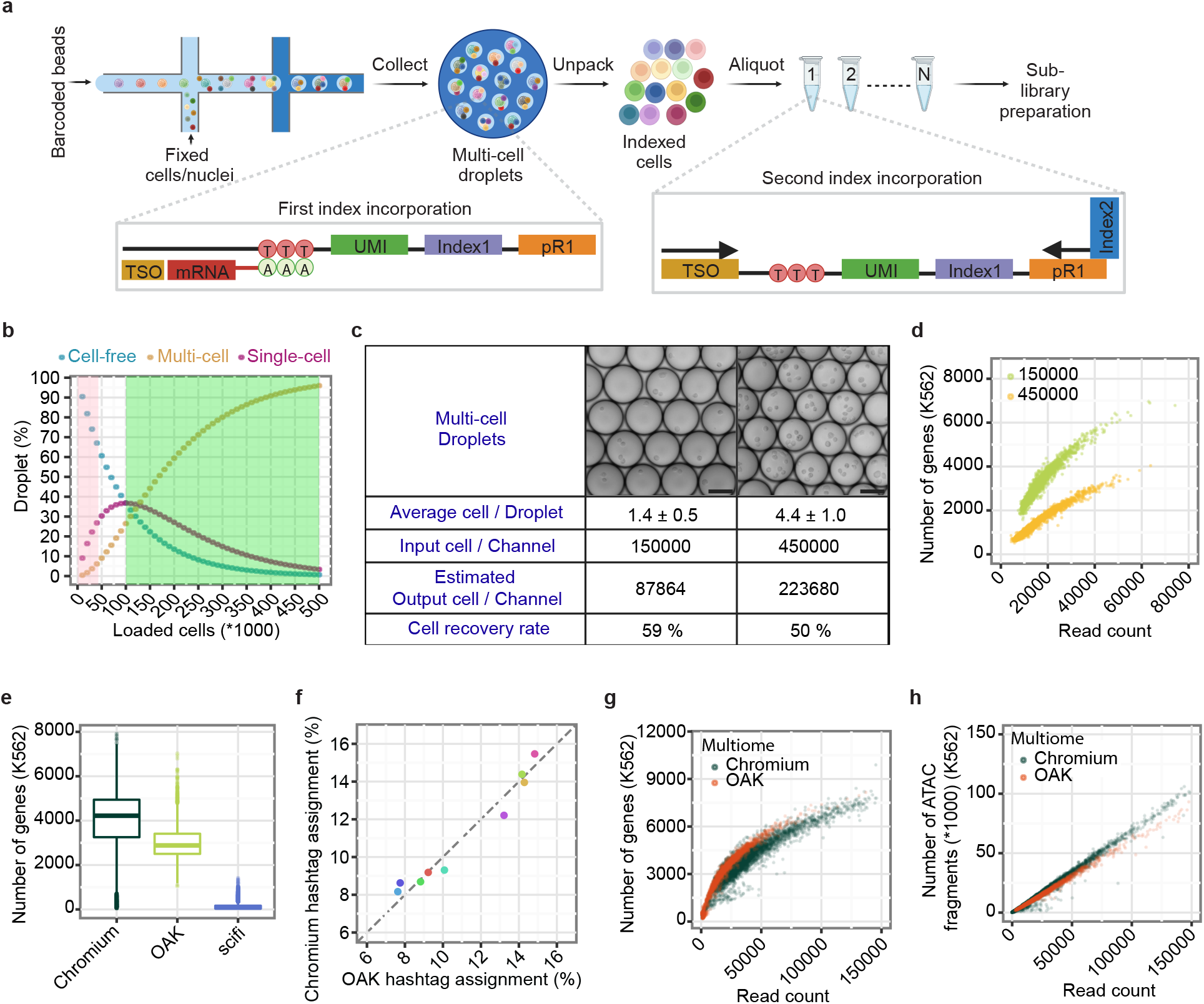
Principle and performance of OAK in single cell profiling of multiple molecular modalities. a, A schematic of the OAK’s scRNA-Seq workflow. During the first indexing step, mRNA molecules hybridize with poly-dT containing bead oligos within droplets in fixed cells or nuclei. Following reverse transcription and emulsion break, fixed cells or nuclei are pooled and re-distributed in individual aliquots where the second index is integrated via a polymerase chain reaction (PCR). TSO: template switch oligo. UMI: unique molecular identifier. pR1: primer binding sequence for TrueSeq Read 1. b, Simulated representation of the percentage of droplets that contain zero (blue), one (magenta), and more than one (yellow) cell, as a function of varying numbers of cells loaded per microfluidic chip channel. Pink and green highlighted areas indicate the range of cell loading in regular 10x Genomics’ Chromium scRNA-Seq and in OAK respectively. c, Droplet images and results of overloading during OAK with different numbers of cells per channel. Scale bars are 75 µm. d, Number of genes detected in K562 cells as a function of the total number of reads per cell. Each data point represents one cell. Green: 150,000 cells loaded, same as in c. Yellow: 450,000 cells loaded, same as in c. e, Number of genes detected in K562 cells with regular Chromium NextGEM 3’ RNA-Seq, OAK scRNA-Seq with 150,000 cells loaded, and scifi-RNA-seq^14^. Boxplots’ center lines represent medians. Box limits denote Q1 (lower) and Q3 (higher) quartiles, and whiskers extend to either 1.5 times the interquartile range (IQR) or to the last data points if they are within these limits. f, Percentage of human bronchial epithelial cells assigned to each sample hashtag (n=9) by the standard Chromium method and OAK for the same sample pool. Each dot corresponds to a different sample hashtag. g, Number of genes detected in K562 cells by joint snRNA-Seq and snATAC-Seq using OAK (red), and standard Chromium (dark green), as a function of the total number of reads per cell. Each data point is a cell. h, Total number of ATAC fragments detected in K562 cells by joint snRNA-Seq and snATAC-Seq using OAK (red) and standard Chromium (dark green), as a function of the total number of reads per cell. Each data point is a cell.

First, to assess the impact of cell overloading, we performed experiments in parallel by loading to a channel on the microfluidic chip 150,000 and 450,000 cells respectively, and compared scRNA-Seq performance at these two cell inputs (Fig. 1c). After sequencing a subset of cells from each experiment, we estimate that 87,864 cells were recovered from the 150,000-cell loading, while 223,680 cells were recovered from the 450,000-cell loading (Fig. 1c). At the same sequencing depth per cell, more genes per cell were detected when 150,000 cells were loaded compared to 450,000 cells (Fig 1d, Extended Data Fig. 1b). The input cells consisted of a 1:1 mixture of a mouse and a human cell line, enabling us to identify collision events when a mouse and a human cell share the same combinatorial indices. When loading 150,000 cells, we found 3.3% cells in the sequencing results to be mix-species multiplets, indicating an overall multiplet rate of 6.6% (Extended Data Fig. 1c), closely aligning with the theoretical expected collision rate (Extended Data Fig. 1a). At the higher loading of 450,000 cells, while we recovered a higher number of cells (Fig. 1c), the overall multiplet rate was 10.6% (Extended Data Fig. 1d). In summary, OAK is flexible to operate with a broad spectrum of loaded cell quantities. The choice on the number of cells to load should be guided by research objectives, balancing between detection sensitivity and cell yield.

We benchmarked OAK to the widely used 10x Genomics’ Chromium NextGEM scRNA-Seq procedure (standard Chromium)^7^, which generates droplet-based RNA-Seq data for up to 10,000 cells per channel on the microfluidic chip. From the 150,000 cells loaded, OAK recovered 87,864 cells per channel (Fig. 1c), a more than eightfold increase in throughput compared to the standard Chromium procedure. With a matched sequencing depth per cell, OAK detected a mean of 3,014 genes per cell for the K562 cell line, while the standard Chromium procedure detected 3,905 genes indicating a mild reduction in sensitivity by OAK (Fig. 1e). Further investigation into the gene detection difference revealed that reduced detection primarily occurred for the lowly expressed genes (Extended Data Fig. 1e). In addition, OAK exhibited a lower percentage of reads that map to mitochondrial DNA (Extended Data Fig. 1f), which is likely attributed to the fixation and permeabilization process that led to partial loss of mitochondria as well as cytoplasmic RNA. This was substantiated by the higher percentage of reads mapping to intronic regions in comparison to the data derived from the standard Chromium procedure (Extended Data Fig. 1g), which also suggests that the permeabilization process could cause an overrepresentation of nuclear mRNA over cytoplasmic mRNA. Overall, a strong correlation between OAK and the standard Chromium method was observed in terms of mean UMI detected across cells for each gene (Spearman correlation coefficient = 0.92, Extended Data Fig. 1h). We compared OAK with scifi-RNA-seq^14^, another combinatorial indexing method utilizing the Chromium system, and observed that OAK exhibited approximately ten times higher sensitivity as measured by number of genes per cell (Fig. 1e).

### Leveraging ultra-high throughput for sample multiplexing

Since OAK enables profiling of hundreds of thousands of cells, it is suitable for ultra-high throughput assays that include many different samples, donors and conditions. Cell hashing with barcoded antibodies is frequently used for multiplexing as it enables pooling of cells from different sources for single-cell profiling^17^. We evaluated antibody hashing within OAK using human bronchial epithelial cells differentiated in transwell plates. We split a sample of antibody stained cells between the OAK and standard Chromium workflow and asked whether cell assignment was comparable. We found that 80% of cells were assigned a hashtag identity in OAK, compared to 81% in the standard Chromium. Furthermore, we found a strong correlation (Pearson correlation coefficient=0.98, Fig. 1f) in the abundance of each hashtag between OAK and standard Chromium. We then clustered cells based on gene expression (Extended Data Fig. 1i). After cell annotation, all expected cell types were present in both data sets, and their proportions correlated between OAK and the standard Chromium (Extended Data Fig. 1j). Therefore, OAK was compatible with the cell hashing approach for sample multiplexing, and did not introduce any biases in cellular composition.

### Flexibility in multimodal single cell profiling

We next investigated whether OAK can perform joint profiling of transcriptome and chromatin accessibility. Since the beads from the Chromium Next GEM Single Cell Multiome kit readily provide barcoding capacity for both mRNA and ATAC fragments, only adjustments in secondary indexing primers were necessary to make OAK compatible with the Chromium multiome workflow (Extended Data Fig. 1k). In order to identify a suitable fixative for joint snRNA-Seq and snATAC-Seq, we evaluated methanol and formaldehyde. Compared to formaldehyde fixation, methanol fixation led to a lower transcription start site (TSS) fragment percentage in the sequencing data (Extended Data Fig. 1l), likely due to methanol’s chromatin denaturing effect. Formaldehyde fixation generated high quality gene expression data (Fig. 1g) and chromatin accessibility data for K562 cells (Fig. 1h). These results underscore OAK’s adaptability in supporting multiple molecular modalities.

### Paired snRNA-Seq and snATAC-Seq for human retinal cells

A common scenario in collecting single-cell data from tissue, is that the most abundant cell types are orders of magnitude higher than the rarest cell types. A couple of examples include recovering neurons from the enteric nervous system, where they represent less than 1% of colon cells^18^, or tuft cells which represent 0.2% of dissociated lung cells^19^. The human retina is another example of a primary tissue with high cellular heterogeneity. Effectively capturing types and states of all cellular subtypes is key to understanding health and disease of the eye; however, the overwhelming presence of rod photoreceptors (around 60% of cells) often impedes the efficient recovery of other cells, such as retinal ganglion cells (less than 1%). Some techniques can be employed for depleting rod cells, however this adds experimental complexity and may unintentionally affect representation of other cell types, for example depletion of bipolar cells^20^. Utilizing a method such as OAK to perform paired snRNA-Seq and snATAC-Seq on retinal samples enables generation of large scale high-resolution data from these precious samples, which are obtained only from careful dissection of post-mortem donations.

We transposed 100,000 fixed peripheral retinal nuclei for overloading (see Methods). We recovered snATAC-Seq data from 42,632 nuclei, and snRNA-Seq data from 46,487 nuclei, with an overlap for 40,691 nuclei. In the snRNA-Seq data we observed a mean of 1,666 genes per cell (Extended Data Fig. 2a). In the snATAC-Seq data we observed the expected fragment distribution pattern (Extended Data Fig. 2b) with a mean of 12,539 fragments per cell and a mean transcription start site (TSS) enrichment of 14.71 (Extended Data Fig. 2c). This compared well to the quality of standard Chromium data generated in parallel for 5,586 nuclei. Standard snRNA-Seq generated a mean of 2,029 genes per cell, whilst the standard snATAC-Seq generated a mean of 14,217 fragments per cell and a mean TSS enrichment score of 12.37.

Using the snRNA-Seq data we clustered and annotated the main cell types of the retina based on known marker genes (Extended Data Fig. 2d, Fig. 2a). With a single donor sample, we obtained thousands of rod, cone, Müller glia, amacrine and bipolar cells, as well as hundreds of horizontal cells, astrocytes and retinal ganglion cells, representing the major cell types of the retina^20–22^. We used the snRNA-Seq annotations with the OAK snATAC-Seq data (Extended Data Fig. 2e) to call open chromatin regions (OCRs) in each cell subtype (Fig. 2b, Extended Data Fig. 2f). We found unique OCR signatures even for the least abundant cell types, including retinal ganglion cells and astrocytes^23^. Peaks called were primarily in intronic and promoter regions as expected (Extended Data Fig. 2g). Looking in more detail at the chromatin peaks in specific cell types we observed differential chromatin accessibility in ARR3 in cone cells (Fig. 2c), and DOK5 in DB5 bipolar cells (Fig. 2d), consistent with previous findings^21^.

**Figure 2.**
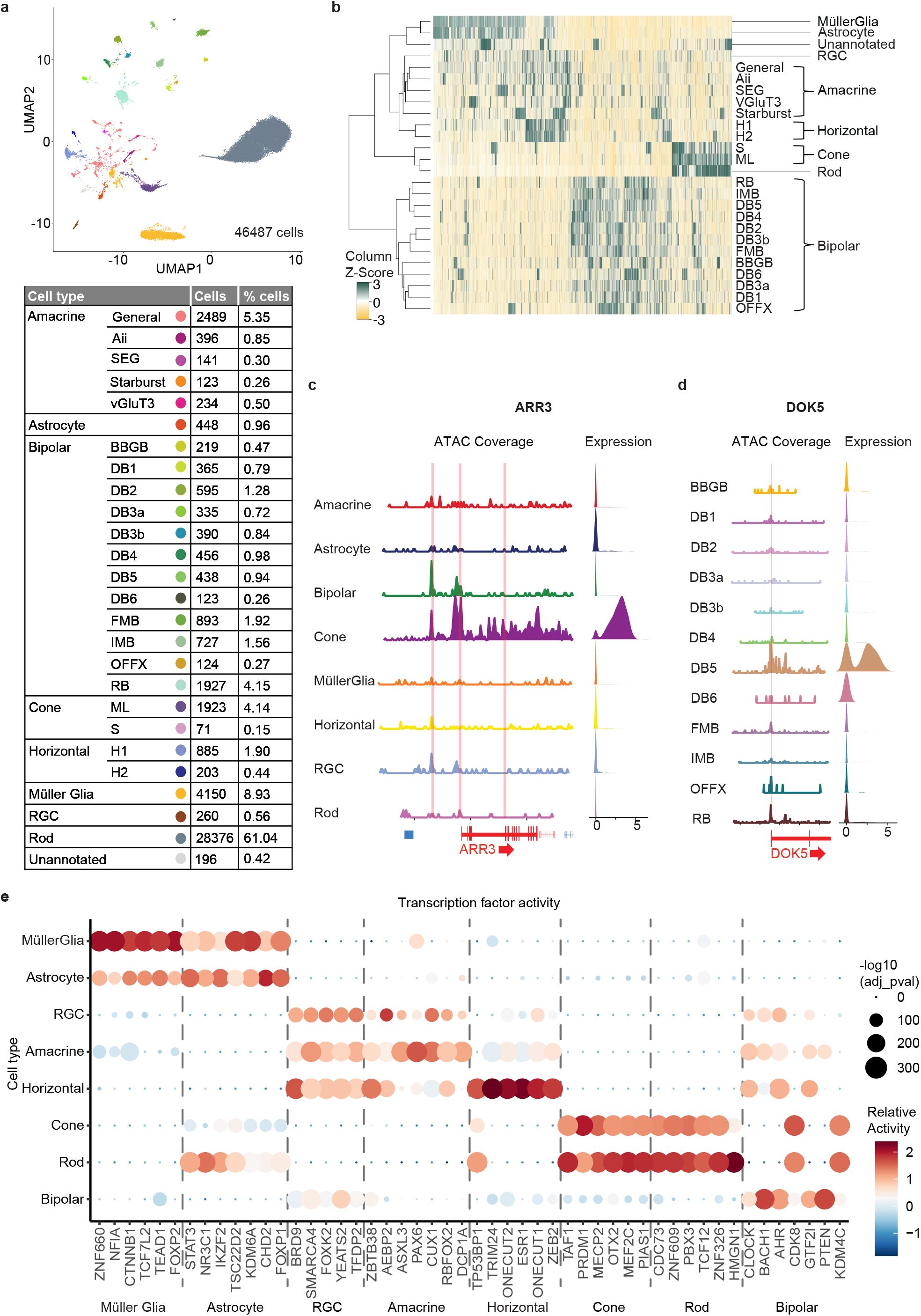
OAK paired snRNA-Seq and snATAC-Seq on the human peripheral retina. a, Uniform Manifold Approximation and Projection (UMAP) of annotated snRNA-Seq data with table of number and percentage of each cell type. The color of the dot for each cell group indicates the position in the UMAP. b, Heatmap displaying detected OCRs in each cell type with FDR <= 0.01 & Log2FC >= 1 using Wilcoxon test for the cell type against a null cell group. c, Chromatin tracks in major cell types for the genomic region (chrX:70253304-70288305) spanning the ARR3 gene, a known cone cell marker, with a Ridge plot (expression values are normalized and log transformed) indicating corresponding gene expression of ARR3 from snRNA-Seq data. d, Chromatin tracks in bipolar cell types for the genomic region (chr20:54455596-54505597) including the TSS of the DOK5 gene, a known marker in DB5 bipolar cells, with a Ridge plot (expression values are normalized and log transformed) indicating corresponding gene expression of DOK5 from snRNA-Seq data. e, Up to seven expressed significant transcription factors by weighted gene activity for each major cell type, as identified by Epiregulon. Transcription factors identified in multiple cell groups are plotted only once. These include transcription factors associated with neuronal cell types: AHR (Horizontal, RGC, Bipolar), BACH1 (RGC, Bipolar) and photoreceptors: CDR73 (Rod, Cone).

Utilizing paired snRNA-Seq and snATAC-Seq data, we identified putative candidates for master regulators in the different cell types using Epiregulon^24^ (Fig. 2e). Epiregulon infers regulatory elements to target genes based on correlated gene expression and chromatin accessibility in clustered cells, matching these elements to known transcription factor binding sites from repositories of public CHIP-Seq data. As a proxy for the strength of the interaction, Epiregulon uses the correlation between transcription factor expression and target gene expression. We plotted the activity for each transcription factor based on the expression of target genes combined with the strength of regulation. We identified elevated BLIMP1/PRDM1 regulation activity in cone cells (Fig. 2e), previously found to be transiently expressed in developing photoreceptors, likely preventing bipolar cell fate^25^. Another example of expected transcription factor activity is of the ONECUT1 and ONECUT2 paralogs activated downstream of PAX6 (Fig. 2e), previously found to be important in the differentiation and maintenance of horizontal cells^26^. Many functional roles of transcription factors in the human retina have been identified by studying early development in analogous animal models or in organoids^27^. Multiomic data generated from post-mitotic cells, as obtained from this retinal sample, offers an intriguing window into ongoing regulation of gene activity decades after initial differentiation events. Obtaining this type of data is especially valuable when considering potential treatments for age-related eye diseases.

### Melanoma resistance to RAF inhibitor belvarafenib

Understanding therapy response and resistance in cancer is crucial for improving treatment outcomes. Belvarafenib is a pan-RAF inhibitor with clinical activity in melanoma^28^. Resistance to belvarafenib arises spontaneously in IPC-298 cells at low frequency^28^. To track emergence of these rare events that could be as infrequent as 0.1%, a substantial cell population is necessary to ensure sufficient representation of the resistant lineages at baseline. By leveraging the high-throughput capabilities of OAK and a lineage tracing technique^29^, we examined transcriptomic response of IPC-298 melanoma cells to a 90-day treatment course with vemurafenib in multiple time points including Day 0, Day 10, Day 20, and Day 90.

We transduced IPC-298 cells with a lentivirus-based library containing 100,000 unique barcode sequences for lineage tracing^29^. A subsample of 1,000 transduced cells, each expected to carry a unique lineage barcode, was expanded. Prior to belvarafenib treatment (Day 0), we collected transcriptomic profiles and lineage barcodes from 144,300 cells (Fig. 3a). The representation of each lineage within the single-cell data displayed a strong correlation with the quantity of reads in bulk sequencing data (Spearman correlation coefficient = 0.93, Extended Data Fig. 3a), confirming accurate lineage recovery with OAK. Furthermore, as the sequenced population of cells increased, the level of correlation between the single cell data and bulk data also increased (Extended Data Fig. 3b), emphasizing the benefit of sampling a high number of cells in systems with such a high lineage diversity.

**Figure 3:**
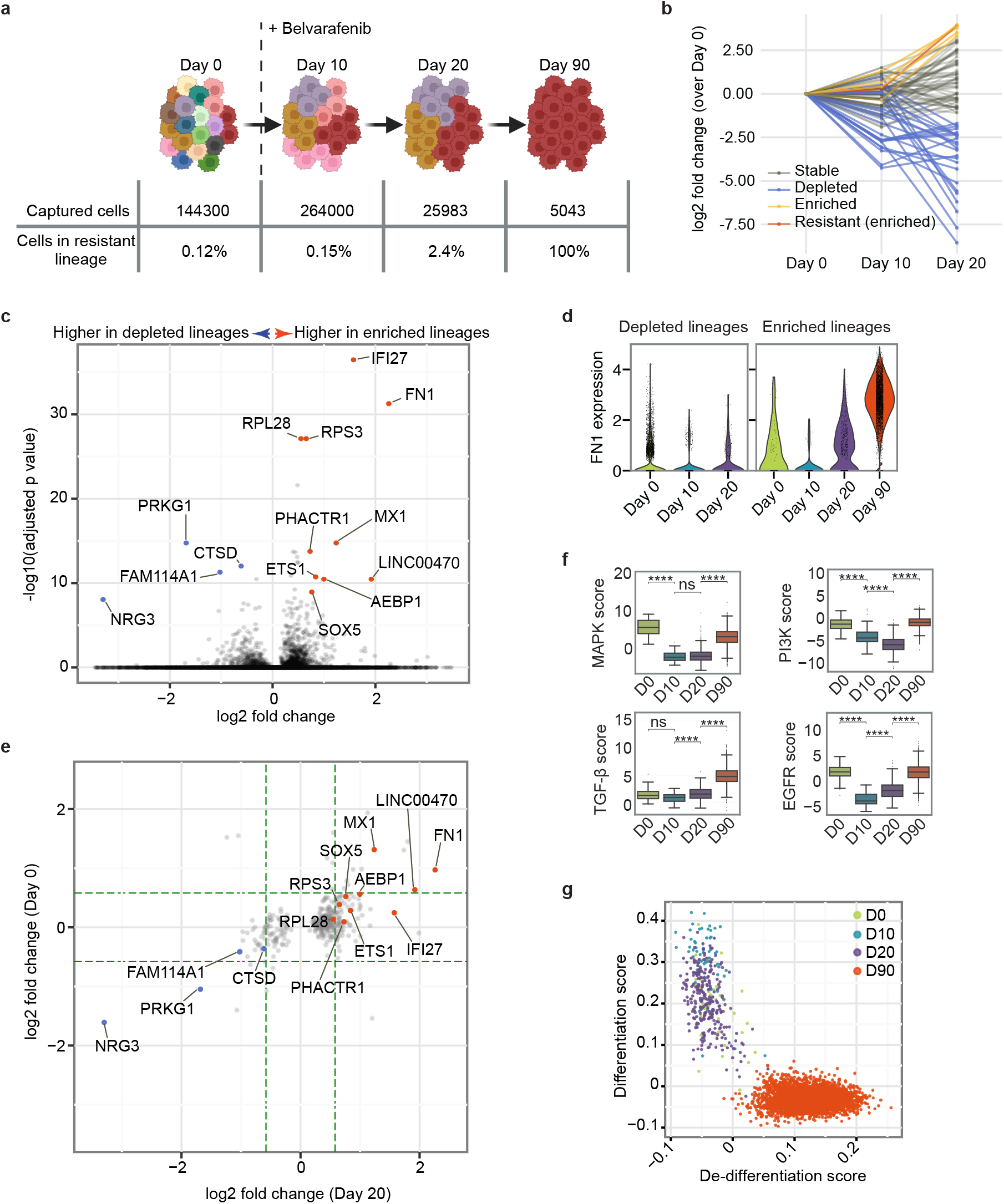
OAK single-cell lineage tracing and transcriptome profiling for melanoma cells during belvarafenib treatment. a, Diagram of the lineage tracing experiment. IPC-298 cells labeled with lineage barcodes were sampled for scRNA-Seq on Days 0, 10, 20, and 90. Belvarafenib treatment commenced following the Day 0 subculture collection. b, Fold change in cell count for each lineage at each time point. Cell counts from Day 0 served as the baseline. Enriched (yellow) includes lineages with over tenfold increase from Day 0 to Day 20, except the resistant lineage. Resistant (enriched) refers to the lineage with over tenfold increase from Day 0 to Day 20, and resistant on Day 90. Stable refers to lineages categorized as neither depleted nor enriched. c, Volcano plot depicting differentially expressed genes identified on Day 20 between depleted and enriched lineages. Genes with adjusted p values lower than 1e-8 and log2 fold changes beyond ±0.5 are labeled. d, Violin plots for FN1 expression level (normalized and log-transformed) at each time point in cells within depleted and enriched lineages. e, Fold changes on Day 0 and Day 20 between the depleted and the enriched lineages. Each data point represents a gene with an adjusted p value <0.05 on Day 20, with specific genes labeled the same as in d. Green dashed lines denote ±1.5-fold changes. f, Scores for PROGENy pathways at each time point for cells within the resistant lineage. ns: adjusted p value > 0.05; ****: adjust p value <=1e-04. Boxplots’ center lines represent medians. Box limits denote Q1 (lower) and Q3 (higher) quartiles, and whiskers extend to either 1.5 times IQR or to the last data points if they are within these limits. g, De-differentiation and differentiation scores for cells within the resistant lineage. Data points are colored based on time points.

Further, we took samples on Day 10 and Day 20 of belvarafenib treatment (Fig. 3a). Five lineages demonstrated over tenfold increase in their relative abundance from Day 0 to Day 20, and therefore were categorized as enriched lineages that are drug tolerant (Fig. 3b). Conversely, 61 lineages, each representing less than 1% of Day 20’s total cells, were defined as depleted lineages (Fig. 3b). After Day 20, as the number of cells continued to drop, we observed the emergence of a belvarafenib-resistant clone among the five enriched lineages (Extended Data Fig. 3c). This clone underwent expansion as a single colony on the plate, and accounted for all of the captured cells on Day 90 (Fig. 3a). Consistent with previous characterization of belvarafenib resistance^28^, this resistant clone only accounted for 0.12% of the cells on Day 0 (Fig. 3a). Thus, cells of that lineage could only be captured with sufficient representation through massive-scale sampling techniques such as OAK. Specifically, for Day 0, we performed stepwise sequencing of sub-libraries, until enough cells from this lineage were recovered for downstream analysis (Extended Data Fig. 3d), and sufficient lineage diversity was achieved (Extended Data Fig. 3e).

To interrogate the transcriptional features associated with drug-tolerance, we computed the marker genes that distinguish the enriched and the depleted lineages on Day 20. We found Fibronectin 1 (FN1) among the overexpressed genes in the enriched lineages (Fig. 3c). Fibronectin-rich extracellular matrix has been shown to provide tolerance for melanoma cells in BRAF inhibition^30^. Moreover, FN1 has been shown to be associated with a mesenchymal phenotype^31^ in melanoma cells. Interestingly, the epithelial mesenchymal transition (EMT) hallmark gene set^32^ emerged as one of the features for Day 90 within the resistant lineage (Extended Data Fig. 3f). In addition, in the depleted lineages FN1 levels remained stable along the course of belvarafenib treatment, while in the enriched lineages the gain of this mesenchymal marker was observed on Day 20 already (Fig. 3d). Moreover, the longitudinal feature of our experiment enabled us to probe for potential pre-existing transcriptional differences between the enriched and depleted lineages. We observed that many of the differentially expressed genes on Day 20 showed differences in expression levels as early as Day 0 (Fig. 3e). This indicates that distinct lineages may possess inherent transcriptional programs for responding to belvarafenib treatment. Furthermore, sustained exposure to belvarafenib led to amplification of selective pre-existing differences, as exemplified by increased fold changes on Day 20 in some of the most differentially expressed genes, such as FN1 and NRG3 (Fig. 3e).

Belvarafenib directly inhibits kinase activity of the RAF kinases, which are responsible for MAPK pathway activation downstream of an oncogenic NRAS mutation in the IPC-298 cells^28^. To assess how the resistant clone adapted to belvarafenib treatment, we specifically compared the activity of the MAPK pathways and several related pathways within the resistant lineage across different time points. We observed an initial downregulation of MAPK, PI3K, and EGFR pathway signatures at early time points, which suggested an initial response to belvarafenib. However, on Day 20 we noticed a rebound of EGFR pathway activity (Fig. 3f). During the same time frame, we observed activation of the transforming growth factor-β (TGF-β) pathway (Fig. 3f), which is a known driver of resistance against MAPK pathway inhibitors in melanoma cells^33^. Furthermore, from Day 20 to Day 90 we observed significant rebound of MAPK and PI3K pathway activities (Fig. 3f), suggesting that reactivation of these pathways may be essential for the establishment of the resistant phenotype.

TGF-β is known to induce EMT^34^ and de-differentiation in melanoma^33,35^. Given a mesenchymal-like state suggested by FN1 upregulation (Fig. 3c), increased TGF-β signaling (Fig. 3f) as early as Day 20, and the enrichment of EMT hallmark genes on Day 90 (Extended Data Fig. 3f), we examined whether the resistant cells switched to a less differentiated state in response to belvarafenib. Despite the initial shift towards a more differentiated melanocyte-like state on Day 10 (Fig. 3g), the resistant cells ultimately reverted to an undifferentiated state (Fig. 3g), resembling the state transitions seen in patient-derived BRAF mutant melanoma cell lines that accompany RAF inhibitor resistance^36,37^.

In summary, our data suggest a progression of transcriptomic alterations along development of belvarafenib resistance. Initial tolerance is associated with activation of EGFR and TGF-β signaling as well as FN1 upregulation. This is followed by MAPK and PI3K pathway reactivation and a shift towards an undifferentiated state, thereby promoting expansion of the resistant cells. OAK’s ultra-high throughput and stepwise sequencing capability render it an exceptionally suitable tool for investigating transcriptomic signatures within rare cell populations that lead to drug resistance in cancer.

## Discussion

OAK combines droplet microfluidics with combinatorial indexing, enabling massive-scale single-cell profiling. Our study underscores its efficacy across diverse experimental designs and modalities, including scRNA-Seq, sample multiplexing, and paired profiling of snRNA-Seq and snATAC-Seq. Moreover, with minor adjustments in the secondary indexing primers and library preparation, broad compatibility can be expected within the full spectrum of applications offered by the Chromium platform, encompassing immune profiling, cell surface protein detection and CRISPR perturbations. In addition, the experimental feature of distributing a large number of cells into smaller aliquots enables sequencing of each sub-library separately. Such stepwise sequencing allows sequencing of a smaller number of cells for quality assessment prior to embarking on large-scale sequencing. Furthermore, sub-libraries provide the opportunity to sequence the number of cells desired for analysis, while preserving unprocessed ones for future data acquisition. Finally, OAK data processing is compatible with analysis pipelines that have been developed for the prevailing commercial Chromium platform. This aspect facilitates a seamless integration of OAK into researchers’ existing data processing workflows.

OAK presents multiple steps for cost savings. First, overloading a single microfluidic channel enables more efficient utilization of costly reagents, including the barcoding beads. Secondly, in contrast to other combinatorial indexing methods^8–11,13,14,38^, OAK avoids a substantial upfront investment in synthesizing plates of indexing oligos or assembling pre-indexed transposome for the ATAC modality - thereby also streamlining benchwork. Thirdly, unlike some overloading methods that identify and discard multi-cell droplets without being able to recover single cells encapsulated within^17,39–41^, OAK is able to resolve single cells in multi-cell droplets, maximizing usage of sequencing data. Ultra-high throughput, extensive versatility for molecular modalities, experimental convenience, and cost efficiency distinguish OAK from alternative technologies in the field.

The single-cell sequencing field is undergoing rapid transformation and growth. Recent examples include innovations like 10x Genomics’ Single Cell Gene Expression High Throughput (HT) and Flex products, both exhibiting superior throughput compared to the regular Chromium products. However, these products are currently unable to perform paired snRNA-Seq and snATAC-Seq. Nevertheless, it is expected that OAK will be adaptable to these evolving platforms, and thereby leveraging improvements in droplet generation technologies to deliver even higher throughput. In this study, our focus rests primarily on validating OAK on the Chromium platform. We expect OAK to be also compatible with other droplet systems with barcoding primers releasable from microspheres, such as the inDrops system^6^ and Hydrop system^16^.

In summary, we developed a new single-cell multiomic profiling method, OAK, which empowers extensive characterization of complex tissues and cellular systems, while maintaining a streamlined and cost-efficient experimental approach. We anticipate that OAK will readily scale with ongoing advances in droplet generation platforms, and will be flexible to accommodate measurement of additional molecular modalities.

## Methods

### Cell culture and single-cell suspension preparation

K562 cells were cultured in Iscove’s Modified Dulbecco’s Medium (IMDM) with 10% fetal bovine serum (FBS). NIH/3T3 cells were cultured in Dulbecco’s Modified Eagle’s Medium (DMEM) with 10% FBS. IPC-298 cells were cultured in RPMI medium with 10% FBS, 2 mM L-glutamine, and 1% penicillin/streptomycin. Cells were incubated at 37°C with 95% Air and 5% CO2. TrypLE™ Express (Thermo Fisher Scientific 12604013) was used to detach adherent cells from culture flasks. Harvested cells were washed twice with phosphate-buffered saline (PBS) with 0.04% Bovine albumin Fraction V (Thermo Fisher Scientific 15260037), and resuspended with PBS to achieve single-cell suspensions.

### Culture and staining of normal human bronchial epithelial cells

Normal human bronchial epithelial cells (Lonza, Epithelix) were differentiated in transwell plates at an air-liquid interface. Cells were dissociated with accutase and then washed twice with PBS/1% Bovine Serum Albumin (BSA). Cells were resuspended in 50 µL PBS/1% BSA and one well of each donor was combined. Nine sample wells were stained with 1 µL of TotalSeq-A antibody, a different hashing antibody was added to each well and incubated at 4°C for 20 minutes. Cells were washed 3x in PBS/1% BSA and then all wells were pooled together. Pooled cells were stained with Sytox Green for 5 minutes at room temperature before sorting for live cells on a Sony SH800S into PBS.

### Retinal tissue nuclei preparation

Human donor eye collection was followed according to a standardized protocol^42^. In brief, the isolated peripheral retinal sample was obtained within a 6-hour post-mortem interval, defined as death-to-preservation time, in collaboration with the Utah Lions Eye Bank. The sample was placed in a cryotube and flash frozen in liquid nitrogen prior to storage at -80°C. The sample was dissociated by douncing ten times in a glass homogenizer in 1 mL ice cold NIM4 buffer (9.9 mL NIM1, 10 µL 100mM DTT, 1 tablet protease inhibitor, 100 µL 10% Triton X-100, 100 µL RNAseIN, 100 µL SUPERasin) and incubated on wet ice for 10 min. Tissue homogenate was centrifuged in at 1500 rpm for 5 min 4°C. Supernatant was removed and 500 µL ice-cold wash buffer (400 µL salt buffer [200 µL 1M Tris pH 7.4, 40 µL 5 M NaCl, 40 µL 5 M NaCl, 60 µL 1 M MgCl_2_, 1.7 mL dH_2_O], 4 µL 100 mM DTT, 40 µL 10% Tween 20, 800 µL 5% RNAse-free BSA, 100 µL RNAse inhibitor, 2.66 mL dH_2_O) was added to the nuclei and the sample pipet mixed five times. The nuclei were passed through a 40 μm filter then counted.

### OAK scRNA-Seq

Single-cell suspension in 400 µl PBS was transferred to a 2 mL round-bottom tube and fixed by adding 1600 µL chilled methanol drop by drop with gentle stirring. Cells were then incubated at - 20°C for 30 min. After fixing, cells were placed on ice for 5 min and then pelleted at 1000G for 5 min at 4°C in a pre-cooled swinging bucket centrifuge. Supernatant was removed and the pellet was resuspended with appropriate volume of resuspension buffer to target 30,000 cells/µl or higher. Resuspension buffer is composed of 3X saline sodium citrate (SSC), 1-2% BSA, 0.2 U/µL Protector RNAse inhibitor, and 1mM DTT. Cells were counted and a desired number of cells (typically 150,000) were loaded per channel. Other reagents were used for loading according to standard 10x Genomics’ Chromium 3’ RNA-Seq protocol. After droplet generation, reactions were transferred to microfuge tubes for reverse transcription at 53°C for 45min.

Immediately after reverse transcription, the droplets were unpacked by adding the recovery agent as described in the standard protocol. After phase separation, the aqueous phase was transferred to a 2 mL microfuge tube. 800 µL 3X SSC was added to the cell suspension. The cells were spun at 650G at 4°C for 5 min. Supernatant was carefully removed. 1 mL 3X SSC was added to the cell pellet with gentle tapping on the tube to dislodge the pellet. Cells were spun again at 650G at 4°C for 5 min. The pellet was resuspended in 215 µL 3X SSC with gentle pipette mixing. A 10 µl solution was used for cell counting to estimate the number of cells per aliquot. The remaining solution was evenly distributed into multiple aliquots (typically 20 aliquots per 150,000 cells loaded to aim for 4,000 cells per aliquot). The aliquots were immediately stored at -80°C until ready for sequencing library preparation.

To prepare sequencing libraries, a desired number of aliquots were heated to 80°C for 5 min to aid release of 1st strand cDNA. Dynabeads^TM^ Silane Viral NA kit (ThermoFisher, 37011D) was used to purify 1st strand cDNA according to manufacturer’s instructions. The cDNA was eluted in 35 µL of the elution buffer. For cDNA amplification PCR, a TSO recognition primer (AAGCAGTGGTATCAACGCAGAGT) and a primer that adds a secondary index (e.g., with barcode underlined: AATGATACGGCGACCACCGAGATCTACAC<u>AACGTGAT</u>ACACTCTTTCCCTACACGACGCTC

TTCCGATCT) were used. For capturing antibody-derived fragments in cell hashing experiments, a single relevant primer can be added, for example, the HTO primer in the case of TotalSeqA hashing. cDNA from multiple aliquots (typically 2-4) can be pooled for sub-library construction by following the standard Chromium protocol, exception in the library PCR where a partial P5 primer (AATGATACGGCGACCACCGAGA) was used alongside an i7 index primer (e.g., with barcode underlined: CAAGCAGAAGACGGCATACGAGAT<u>CGCATGTTAC</u>GTGACTGGAGTTCAGACGTGT.

After QC, sub-libraries with unique secondary index (i5) and i7 index were pooled for sequencing on Illumina platforms, with 28 cycles for Read 1, 10 cycles for i7 index, 8 cycles for i5 index, and 90 cycles for Read 2. Targeted sequencing depth was 20,000 read pairs per cell. Cell counting at the aliquoting step was used to estimate the number of cells expected to recover.

### OAK paired snRNA-Seq and snATAC-Seq

Nuclei were centrifuged at 500G in a 2 mL round-bottom microfuge tube. After removing the supernatant, the pellet was resuspended in a fixation solution of 1 mL calcium-free PBS with 0.3% formaldehyde. Nuclei in fixation solution were placed on ice for 10 min and then centrifuged for 5 min at 500G at 4°C. After supernatant was removed, 1.5 mL wash buffer was added. The wash buffer was 10 mM Tris-HCl (pH 7.4), 10 mM NaCl, 3 mM MgCl2, 1% BSA, 0.1% Tween-20, 1 mM DTT, 1 U/μL RNase inhibitor in nuclease-free water. After 5 min 500G at 4°C, the supernatant was removed and the nuclei were resuspended in the appropriate volume of nuclei resuspension buffer to target 2,400 nuclei/μL or more. The nuclei resuspension buffer was 1X Nuclei Buffer (from a 20X stock, 10x Genomics, PN2000207), 1mM DTT, and 1 U/μL RNase inhibitor in nuclease-free water.

After fixation, we typically transpose 75,000-200,000 nuclei, with an expected cell recovery rate of 38%-42%. This recovery rate may be dependent on sample type and quality. We found that TDE1 enzyme (Illumina Tagment DNA Enzyme and Buffer Small Kit, 20034197) could be used for the additional tagmentation reactions required to support processing large numbers of nuclei. Each transposition reaction was composed of 12,000 nuclei in 5 μL 1X nuclei buffer, 3 μL TDE1 enzyme, and 7 μL ATAC Buffer B (10x Genomics, PN 2000193). Reactions were incubated at 37°C for 1 hr. All transposition reactions were combined to a 2 mL round-bottom microfuge tube and spun at 500 G in a pre-cooled centrifuge at 4°C for 5 min. Supernatant was removed, leaving transposed nuclei in 15 μL of solution which was used for loading 1 channel. Other reagents were used for loading according to standard 10x Genomics’ Chromium Next GEM Single Cell Multiome protocol. After GEM generation, barcoding, and quenching according to the standard protocol, 125 μL recovery agent (10x Genomics, PN 220016) was added to break the emulsion. The aqueous layer was carefully transferred to a 2 mL round-bottom microfuge tube. 800 µL 3X SSC was added to the cell suspension. The cells were spun at 650G at 4°C for 5 min. Supernatant was carefully removed. 1 mL 3X SSC was added to the cell pellet with gentle tapping on the tube to dislodge the pellet. Cells were spun again at 650G at 4°C for 5 min. The pellet was resuspended in 215 µL 3X SSC with gentle pipette mixing. A 10 µl solution was used for cell counting to estimate the number of cells per aliquot. The remaining solution was evenly distributed into multiple aliquots (typically 20 aliquots per 150,000 cells loaded to aim for 4,000 cells per aliquot). The aliquots were immediately stored at -80°C until ready for sequencing library preparation.

To prepare sequencing libraries, a desired number of aliquots were heated to 80°C for 5 min to aid release of 1st strand cDNA and ATAC fragments. Dynabeads^TM^ Silane Viral NA kit (ThermoFisher, 37011D) was used to purify 1st strand cDNA and ATAC fragments according to manufacturer’s instructions. The resulting products were pre-amplified in a 100 μL reaction using 10 cycles with 4 μL pre-amp primers (10x Genomics PN 20002714) and 50 μL of NEBNext High-Fidelity 2X PCR Master Mix (NEB, M0541S). Reactions were cleaned with 1.6X SPRI and eluted in 40 μL EB. 10 μL of the product was used for constructing snATAC-Seq libraries. One PCR reaction was set up for each aliquot, with 0.6 μL of 100 μM partial P5 primer, 50 μL NEBNext High-Fidelity 2X PCR Master Mix (NEB, M0541S), 36.9 μL nuclease-free water and 2.5 μL of 10 μM sample index N (10x Genomics, PN 1000212). The PCR program was 98°C 30s, n cycles [98°C 10s, 67°C 30s, 72°C 20s], 72°C 2 min, held at 4°C. N is typically recommended cycles for standard Chromium protocol for the number of cells. Extra cycles can be added if ATAC library yield is low. A double-sided size selection was performed as instructed in the standard Chromium protocol. cDNA library amplification, sequencing library construction and sequencer operation were conducted in the same way as described in OAK scRNA-Seq.

For snATAC-Seq libraries, sub-libraries with unique i7 index were pooled for sequencing on Illumina platforms. Targeted sequencing depth was 25,000 read pairs per cell, with 50 cycles for Read 1, 8 cycles for i7 index, 24 cycles for i5 index, and 49 cycles for Read 2. Cell counting at the aliquoting step was used to estimate the number of cells expected to recover.

### Sequencing read processing

Illumina Miseq, Nextseq 2000, and NovaSeq 6000 were used for sequencing. Raw sequencing data was demultiplexed by Illumina’s Bcl2Fastq software to resolve reads per OAK sub-libraries. Fastq files for each sub-library were processed with Cell Ranger software v6 (single-cell RNAseq) or Cell Ranger ARC software v2 (paired snRNA-Seq and snATAC-Seq) to generate gene and chromatin fragment counts.

### Simulation for cell distribution in droplets

The percentage of having k cells in a droplet is approximated by p(k, λ)=e(-λ)*λk/(k)! based on Poisson distribution, where λ is the loading rate approximated as the number of loaded cells divided by the number of generated droplets.

### Multiplet rate theoretical estimation

The expected number of events when more than one cell share the same combinatorial barcodes is N-D+D*[(D-1)/D]N based on the closed form solution for expected number of collisions in the birthday paradox^11^, where N is the number of cells loaded and D is the total number of barcode combinations. We used 100,000 as the number of droplets generated per channel on the Chromium microfluidic chip. Hence D was calculated as 100000*n_aliquot, where n_aliquot is the number of aliquots generated.

### Species-mixing experiment and multiplet rate estimate

In the species-mixing experiment when 150,000 cells (Fig. 1c) were loaded in one channel of the Chromium chip, all cells were unpacked into 12 aliquots. In the experiment when 450,000 cells (Fig. 1c) were loaded in one channel, all cells were unpacked into 40 aliquots. In each experiment, one aliquot was processed to generate a sequencing library. Reads were mapped to a hybrid Human-Mouse reference genome that consists of GRCh38 and mm10. Cells were classified into observed multiplets (human+mouse), mouse cells, and human cells by Cell Ranger software v6. Since the input cells consisted of a 1:1 mixture of a mouse and a human cell line, true multiplet rate was estimated as (observed multiplet rate)*2 to include those inferred human+human and mouse+mouse multiplets.

### Hashtag assignment and cell annotation in human bronchial epithelial cells

The Cell Ranger package was used to determine hashtag assignment rate for the human bronchial epithelial cell experiment using a matching antibody-derived tag and gene expression library for each of four (out of 22) OAK aliquots, and for standard comparison data generated in parallel. We imported and merged the data from the multiple OAK aliquots in Seurat, then integrated the OAK and standard scRNA-Seq data with Harmony^43^ prior to clustering using the Seurat FindClusters function (resolution = 0.6) and assigning cluster identity (Club/Goblet, Basal, Ciliated, Basal cycling, Neuroendocrine or Unknown) based on gene scores for known markers.

### Retinal paired snRNA-Seq and snATAC-Seq data analysis

snATAC-Seq and gene expression data from each sub-library were combined using cell ranger-arc aggr with -normalize=none. Gene expression data was first imported into a Seurat (Version 4.3.0.1) object for assessment of snRNA-Seq quality and comprehensive annotation. Cells with >200 genes and <10000 genes were retained. Cells were clustered, then marker gene scores were used to validate assignment of clusters to the major known cell types. For cones, horizontal, amacrine and bipolar cells, further sub-clustering was performed prior to annotation and propagation into the master Seurat object. The snRNA-Seq annotations were added as metadata into an ArchR project containing both snATAC-Seq and snRNA-Seq data, based on cell barcodes. For further analysis of snATAC-Seq data, only cells with data passing filters from both modalities were kept.

In ArchR (Version 1.0.2), hg38 was used as the reference genome and barcodes were filtered for TSS enrichment >4 and nFrags >1000. Peaks of open chromatin were identified by using ArchR tools. First addReproduciblePeakSet was utilized with MACSr, which uses the MACS3 algorithm for peak-calling ^44^, on the snRNA-Seq annotation, excluding chrMT and ’chrY, followed by addPeakMatrix. Marker peaks were called using this matrix for each cell type with getMarkerFeatures with options bias = c(”TSSEnrichment”, “log10(nFrags)”) and testMethod=”wilcoxon”. Peaks called were filtered for FDR cutOff of 0.01 and Log2FC >= 1. The plotMarkerHeatmap function was used to plot a heatmap of the markers with FDR <= 0.001 and Log2FC >= 1. The plotBrowserTrack function in ArchR was used to plot example chromatin tracks and peaks for each annotated cell group.

Epiregulon infers regulatory elements to target genes based on correlated gene expression and chromatin accessibility in clustered cells, matching these elements to known transcription factor binding sites from repositories of public ChIP-Seq data. In Epiregulon (Version 1.0.34) we extracted the normalized gene expression counts and peak matrices from the ArchR project, removing unannotated cells, and calculated the peak to gene expression linkages using the LSI snRNA-Seq and snATAC-Seq combined dimensions from ArchR. We annotated the linkages as regulons with known motifs from the human ChIP-Atlas and Encode databases. We further pruned the regulons by setting a correlation test cutoff for for all components (peaks, gene expression and TFs in the same cells) using the following parameters to the pruneRegulon function (test = “chi.sq”, prune_value = “pval”, regulon_cutoff = 0.05 and defined clusters by the major cell types). We used the addWeights function to add an estimate and multiplier for the strength of regulation, using the parameters tf_re.merge = FALSE, method = “corr”. We calculated a score for each regulon using calculateActivity to combine weights with activity of linked genes (mode = “weight”, method = “weightedMean”, exp_assay = “normalizedCounts”, normalize = FALSE). To find the transcription factors with differential regulation activity associated with each major cell type we used the findDifferentialActivity function with parameters, pval.type = “some”, direction = “up” and test.type = “t”). We filtered these by significant transcription factors with an FDR cutoff of 0.05 and a logFC cutoff of 0.1. With this list in hand, we added information on the proportion of cells that have expression of the identified transcription factor in the significant group. We filtered the transcription factors to those that have >30% expression in the associated cell type. We took the top ten transcription factors with the highest calculated activity in each cell type and ordered them by the proportion of cells expressing within the cell type. We then used the plotBubble function to plot the top seven transcription factors with the highest calculated activity for each cell type.

### TraCe-seq cell preparation and scRNA-Seq

TraCe-seq barcode lentivirus was produced, and cells were infected and sorted as described in the previous publication ^29^. After sorting, 1,000 IPC-298 cells were used to form the starting population. This population was expanded for 17 doublings. A subculture was used for the Day 0 experiment, while the rest of cells were treated with 10 µM belvarafenib. Medium containing belvarafenib was replenished twice a week. Subcultures of cells were taken for OAK on Day 0 and Day 10 as lineage diversity was highest before and early in treatment. For Day 0, 2 channels on the Chromium chip were loaded, each with 138,000 cells. 39 aliquots were generated, each contained 3700 cells. 20 aliquots were processed into sub-libraries and sequenced. For Day 10, 3 channels were loaded, each with 180,000 cells. 44 aliquots were generated, each contained 6000 cells. 12 aliquots were processed into sub-libraries and sequenced. The remaining aliquots were stocked in -80°C for potential future data acquisition. Standard Chromium scRNAseq was performed according to manufacturer’s instructions for cells collected on Day 20 and Day 90 of treatment as lineage diversity dropped.

### TraCe-seq single-cell lineage barcode library generation

OAK indexed cDNA libraries from multiple aliquots can be pooled for the generation of lineage barcode libraries. Typically, 7.5 µl cDNA from each aliquot is used and 2 aliquots were pooled for 1 reaction. A semi-nested PCR strategy was used to ensure the specificity of the resulting lineage barcode library. In the first round of PCR, the partial P5 primer and GPF_F1_outer primer (GTGCACTTAGTAAGGACCCAAACG) were used. In the second round of PCR, the partial P5 primer and an i7 indexed GFP_F2_inner primer (e.g., with index underlined:CAAGCAGAAGACGGCATACGAGAT<u>CCGCGGTT</u>GTGACTGGAGTTCAGACGTGT GCTCTTCCGATCTGATAACCCTCGGGATGGATGAACTG) were used.

### TraCe-seq bulk lineage library generation

Cells from Day 0 were used to amplify the lineage transcripts. The reverse transcription mix was composed of 5 µL Maxima H minus Reverse Transcriptase (Thermo Fisher Scientific EP0753), 20 µL 5X RT buffer, 5 µL dNTP (10 mM each), 1.5 µL TraCe_libABC_end_RT primer (GTGGATCCACCGAACGCAACGCAC, 100 µM), 1.5 µL Protector RNase Inhibitor (Sigma PN 3335399001), 5 µl methanol fixed cells, and 62 µl water. The reaction was incubated at 50°C for 30 min, followed by 85°C for 5 min, and held at 4°C briefly. The product was subsequently amplified by PCR with P5 indexed primer (e.g., with index underlined: AATGATACGGCGACCACCGAGATCTACAC<u>GATATCGA</u>CGAACGCAACGCACGCACACT) and i7 indexed GFP_F2_inner primer. The SPRISelect beads were used to perform a 0.6X-1.6X double sided size selection for the PCR product.

### Drug response curve generation

Cells were seeded at 2,000 cells per well in 96-well plate, and were treated with belvarafenib 24 hours after seeding. Cells were treated with a 9-point titration (1:3) and DMSO control using the HP D300 drug dispenser. Cell growth was assessed using CellTiter-Glo Luminescent Cell Viability Assays (Promega G7570), and luminescence was read by a 2104 EnVision Multilabel Plate Reader (PerkinElmer) five days after treatment. All cell viability data was collected and calculated for 4 replicates per condition. Data from the DMSO control was set to 100%. Nonlinear regression curves were generated by GraphPad Prism to fit the viability data.

### TraCe-seq data analysis

Cells were assigned to a lineage when the UMI count for one lineage barcode was at least two-fold higher than the other ones detected in the given cell. Single-cell gene expression matrix was analyzed with Scanpy^45^. Gene set enrichment for MSigDB’s hallmark sets^32^ was performed with decoupleR^46^. MAPK, EGFR, PI3K, and TGF-β pathway scores were generated with PROGENy^47^. P values were calculated using the Mann-Whitney-Wilcoxon test (two-sided) with Bonferroni adjustment. Genes representing differentiation and dedifferentiation states were based on an established melanoma four-stage differentiation model^48^. The melanocytic, transitory-melanocytic, transitory, and neural crest-like-transitory signatures were grouped as the differentiation signature. The undifferentiated, undifferentiated-neural crest-like, and neural crest-like signatures were grouped as the de-differentiation signature. The signature scores were generated by Scanpy’s tl.score_genes function.

Schematics used in this manuscript were created with BioRender.com.

### Data and code availability

Sequencing data and code to be made available upon request.

## Acknowledgements

We thank the donor and the donor’s family who contributed the retinal sample for this study. We acknowledge the single-cell sequencing community at Genentech for collaborations, discussions, and sequencing support, including Ahmet Kurdoglu, Qixin Bei, Manching Ku, Jie Liu, Daniel Le, Ashley Byrne, William Stephenson, Vasu Kameswaran, Bence Daniel, Jay Leone, Shiqi Russell Xie, Diana Wu, Katie Geiger-Schuller, Ana Meireles, Kristel Dorighi, Aviv Regev, Xiaosai Yao, David Garfield, Luz Orozco, Natalie Fox, Jack Kamm, Lyndsay Murrow and Jennie Lill.

## Contributions

B.W. conceived and developed OAK. B.W. and H.M.B. optimized the methods. B.W., H.M.B., X.Y., and S.D. designed the study. M.D., L.A.O., I.K.K. provided the retinal sample. H.M.B. and A.S. conducted the experiments on the retinal sample and H.M.B. analyzed the retinal data. H.M.B., E.V.N., C.E. and C.E. conducted the experiments on the *in vitro* differentiation bronchial samples, H.M.B. and S.D. analyzed the data. B.W. and X.Y. performed the melanoma cell lineage tracing experiments and B.W. analyzed the sequencing data. C.C. and J.L. generated sequencing libraries. M.S., J.M.L., A.X-M., N.P., and Y.L. performed sequencing. B.W., H.M.B., X.Y., S.D., and Z.M. wrote the manuscript.

## Ethics declarations

B.W., H.M.B., X.Y., A.S., C.E., C.E., E.V.N., H.C., C.C., J.L., M.S., J.M.L., A.X.M., N.P., Y.L., Z.M. and S.D. are currently employees and shareholders of Genentech, a member of the Roche Group.

## Figure Legends

**Extended Data Figure 1: OAK assay performance and compatibility with multiple molecular modalities.**

a, Collision rate corresponding to each number of total aliquots generated per channel, with 150,000 cells (green) or 450,000 cells (yellow) loaded per channel.

b, Number of genes detected in NIH/3T3 cells as a function of the total number of reads. Each data point represents one cell. 150,000 cells (green) and 450,000 cells (yellow) were loaded respectively, same as in Fig. 1c.

c-d, Total number of mouse UMIs and human UMIs detected in each combinatorial index, when 150,000 cells (in c) or 450,000 cells (in d) were loaded per channel. Combinatorial indexes were classified as human cells (red), mouse cells (blue), or observed multiplets (green).

e, Percentage of genes expressed in K562 cells detected by the standard Chromium method that were recovered by OAK in each of the expression percentile bins set by expression levels. f, Percentage of reads mapping to the mitochondrial genome in K562 data collected by the standard Chromium 3’ RNA-Seq and OAK’s scRNA-Seq.

g, Percentage of reads mapping to intronic regions in K562 data collected by the standard Chromium 3’ RNA-Seq method and OAK’s scRNA-Seq.

h, Number of UMIs per gene in K562 cells detected by the standard Chromium 3’ RNA-Seq and OAK’s scRNA-Seq. Each data point is a gene.

i, UMAPs for in vitro differentiated bronchial airway cells profiled by OAK and standard Chromium method. For comparative visualization, data from both methods was integrated with Harmony^43^ and the clusters were annotated using markers for major cell types.

j, Percentage of cells for each cell type annotated within the OAK and the standard Chromium dataset. Basal (cycl.): Basal cycling cells. NE: Neuroendocrine cells.

k, Schematic diagram of the experimental procedure for OAK’s multiome (joint snRNA-Seq and snATAC-Seq). Building on Fig 1.a, cells or nuclei are fixed and then transposed to generate ATAC fragments before droplet generation. First indexing occurs in each droplet by hybridizing the poly-dT containing bead oligos with mRNA molecules, as well as by ligating spacer sequence containing bead oligos with transposed chromatin fragments through a bridge oligo. Second indexing occurs in each aliquot by PCR with a pair of barcoded primers for the cDNA and a pair of barcoded primers for the chromatin fragments.

l, Percentage of fragments in K562 data that overlap transcription start sites (TSS). Chromium: standard Chromium multiome (joint snRNA-Seq and snATAC-Seq); OAK_FA: OAK-multiome (joint snRNA-Seq and snATAC-Seq) with formaldehyde as fixative; OAK_MeOH: OAK-multiome with methanol as fixative.

**Extended Data Figure 2: OAK data analysis of paired snRNA-Seq and snATAC-Seq on human peripheral retina.**

a, Number of genes detected in human retinal cells as a function of the total number of reads per cell. Each data point represents one cell. The dotted line represents the mean value.

b, The snATAC-Seq fragment size distribution for all cells for 535.36 million fragments.

c, Density plot with histogram of TSS enrichment and number of unique fragments for each cell in snATAC-Seq data. Vertical line represents the mean number of fragments per cell and the horizontal dotted line represents the mean TSS score per cell.

d, UMAP of unannotated RNA clustering.

e, UMAPs of snATAC-Seq data showing cluster assignment (left) and transfer of snRNA-Seq annotations (right) based on cell barcodes.

f, Heatmap of snATAC-Seq peaks clustered for each bipolar cell type.

g, Number of snATAC-Seq peaks called for each cell type in relation to annotated intronic, promoter, exonic and distal gene regions.

**Extended Data Figure 3: OAK enables lineage tracing, identifying temporal gene signatures within a resistant melanoma cell lineage.**

a, Correlation of lineage abundance between measurements by the number of cells in OAK data and by the number of reads in bulk sequencing data on Day 0. Each data point represents a lineage.

b, Spearman correlation coefficients between lineage abundance measured by OAK and by bulk sequencing against varying numbers of cells sequenced for Day 0. Each data point’s cells were generated by randomly downsampling from the total number of sequenced cells (74,000). Each downsampling was iterated 10 times. Boxplots’ center lines represent medians. Box limits denote Q1 (lower) and Q3 (higher) quartiles, and whiskers extend to either 1.5 times the IQR or to the last data points if they are within these limits.

c, Cellular viability of parental cells and the belvarafenib-resistant clone treated by increasing concentrations of belvarafenib. Data are mean ± s.e.m, with 4 replicates per concentration per group of cells.

d, The cumulative number of cells detected within the resistant lineage on Day 0 when a varying number of sub-libraries were sequenced.

e, The cumulative percentage of lineages recovered on Day 0 when a different subset of sub-libraries (X-axis, bottom) were sequenced. The total number of cells sequenced in the corresponding number of sub-libraries are indicated on the top.

f, Heatmap for gene set scores at each time point for cells within the resistant lineage. At each time point, hallmark gene sets with mean score changes >0.3 and adjusted p values < 0.01 were displayed.

