## Supplementary figures and images for "Overloading And unpacKing (OAK) - droplet-based combinatorial indexing for ultra-high throughput single-cell multiomic profiling"

### Extended Data Figure 1

**Extended Data Figure 1**

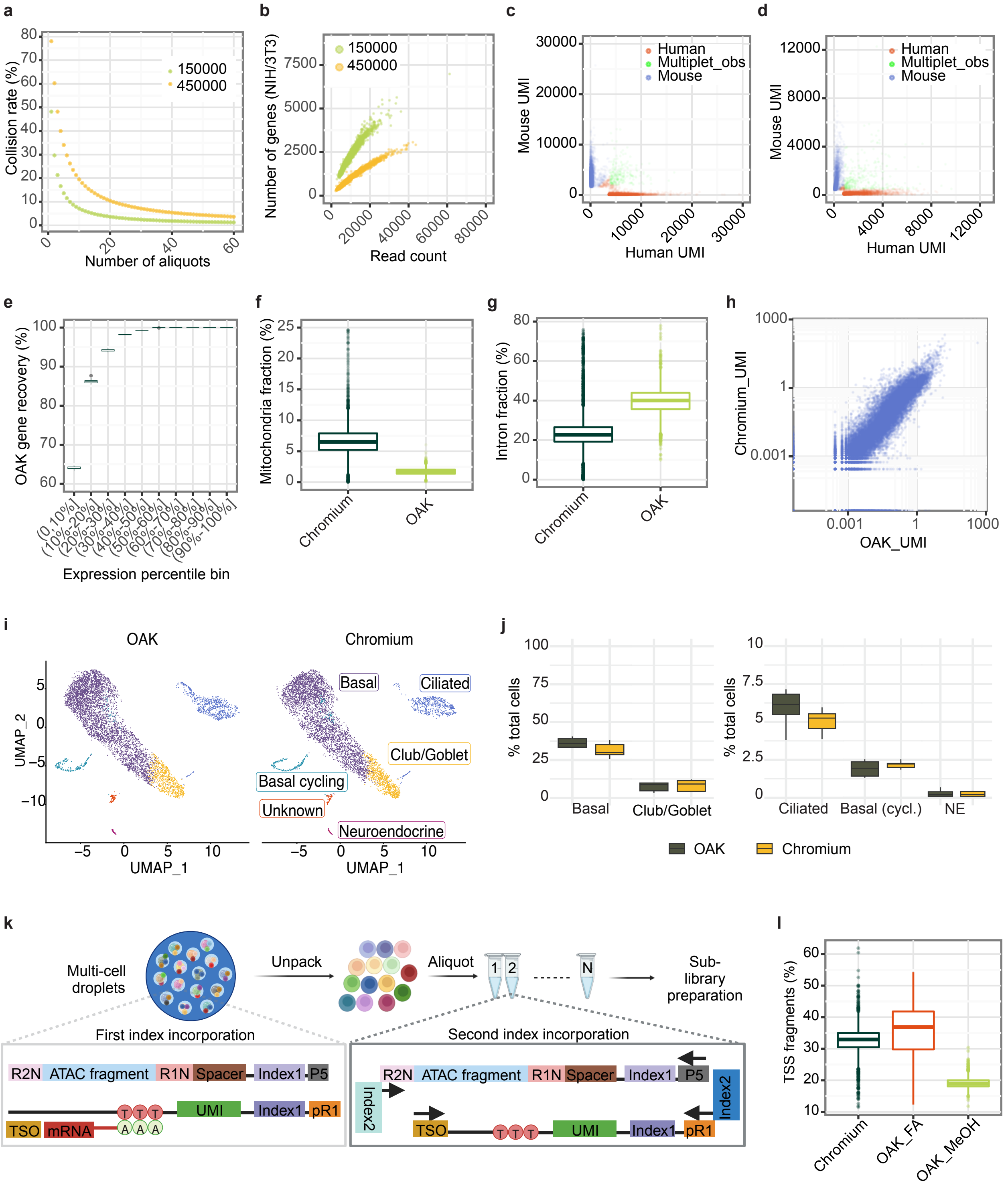

### Extended Data Figure 2

**Extended Data Figure 2**

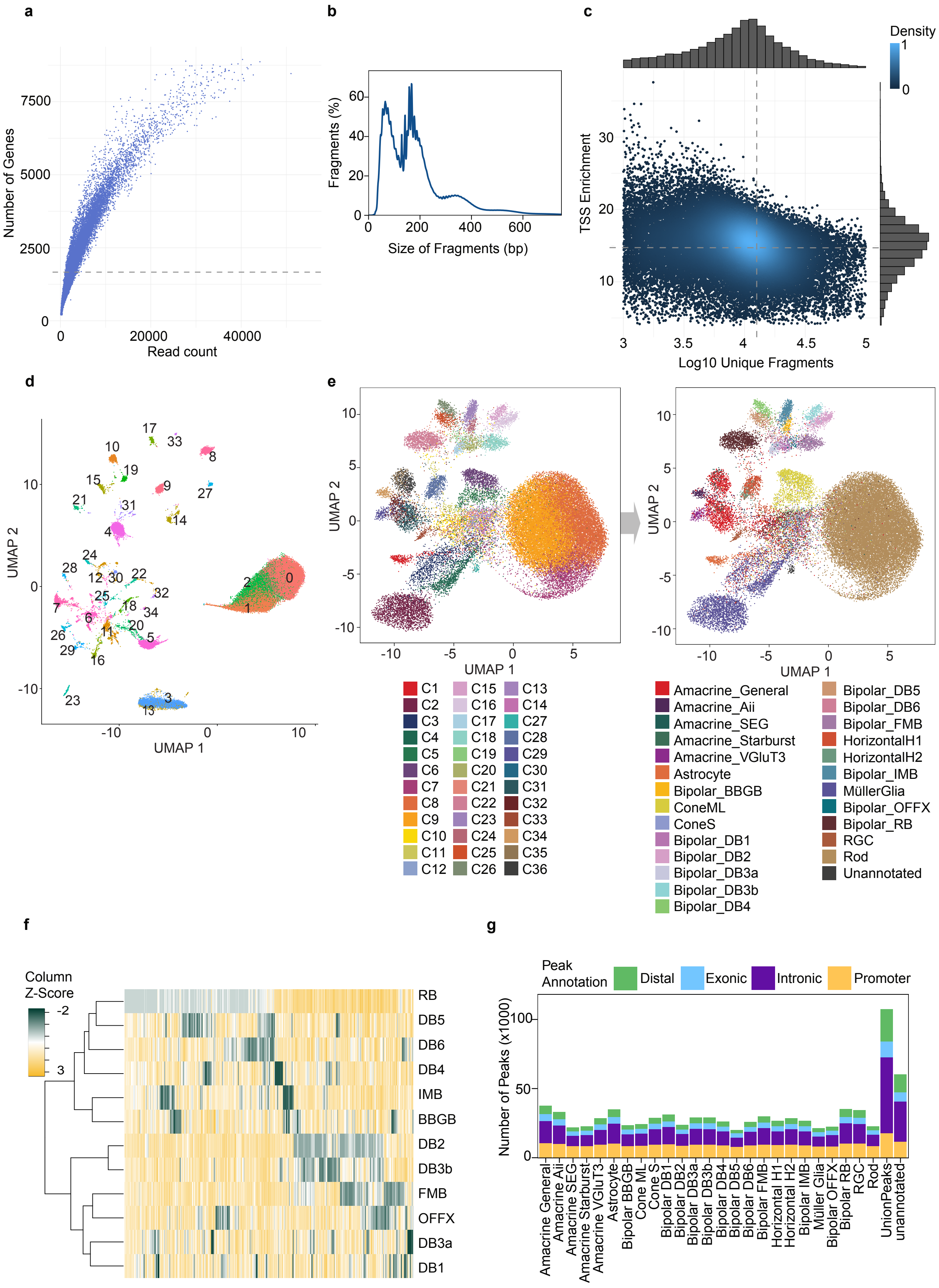

### Extended Data Figure 3

**Extended Data Figure 3**

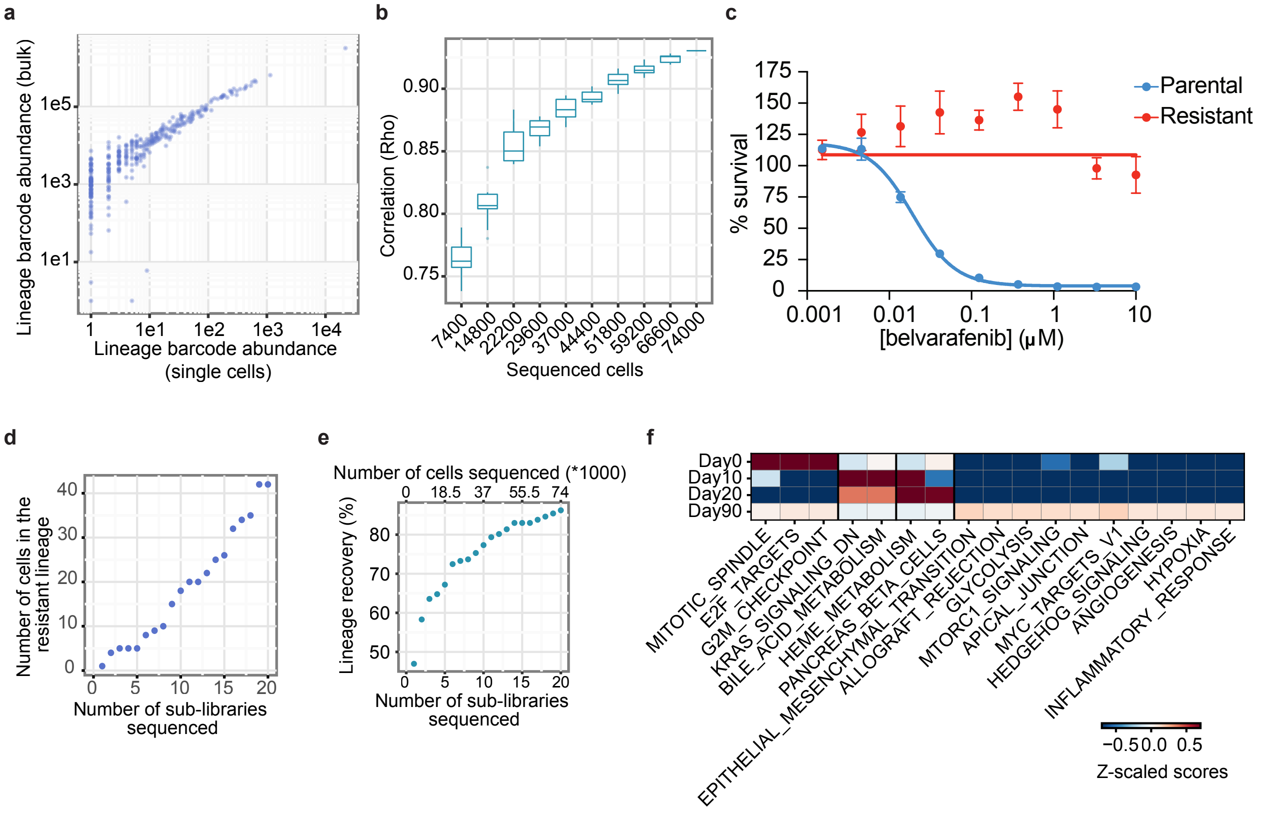
